# Isolation by distance and metapopulation structure in Mozambican populations of the soft tick *Ornithodoros phacochoerus*, a vector of African Swine Fever virus

**DOI:** 10.1101/2025.10.07.680904

**Authors:** Florian Taraveau, Carla Giupponi, David Bru, Jacinto Chapala, Antónia De Andrade, Maxime Duhayon, Alberto Francisco, Karine Huber, Mélanie Jeanneau, Elsa Rudo Pires Lameira, Thomas Pollet, Carlos João Quembo, Hélène Jourdan-Pineau

**Author notes:** Equal contribution (Florian Taraveau, Carla Giupponi).

## Abstract

Ticks of the species *Ornithodoros phacochoerus* are endophilic soft ticks which infest warthog burrows. Like other *Ornithodoros* of the *O. moubata* species complex, *O. phacochoerus* is a vector of the African swine fever virus and participates in the maintenance of the virus among warthogs in the sylvatic cycle in Southern and Eastern African countries. In this study, the population genetic structure of *O. phacochoerus* was investigated using fifteen microsatellite markers. After sampling campaigns that took place between 2020 and 2024, we analyzed 679 ticks from 22 sites (warthog burrows and resting sites) from Coutada 9 Game Reserve and 368 ticks from 10 sites from Gorongosa National Park in Mozambique. Gorongosa National Park and Coutada 9 Game Reserve appeared to be completely isolated from each other in terms of gene flow. In the two conservation areas, the observed genetic variation followed a model of isolation by distance with a structure at the level of the sampling sites. Effective population sizes within sampling sites were quite low and signals of bottlenecks were found in various sites in the two conservation areas. These results suggested a metapopulation system with gene flow occurring among neighbouring populations, likely due to tick transfer by their hosts. Such migration pattern participates in ASF virus dissemination by soft ticks at a small geographic scale, highlighting the importance of maintaining buffer zones around conservation areas to prevent vector spillover from the sylvatic cycle to the domestic cycle.

**Author summary:** African swine fever is a viral disease that is often fatal to domestic pigs and causes major economic losses worldwide. In Southern Africa, the virus is maintained in a cycle involving warthogs, which are resistant to the virus, and soft ticks living in their burrows. For decades, we have known that these ticks can transmit and maintain the virus in both wild and domestic settings, but not whether they may be involved in spillover of the virus between the two compartments. In this context, we sought to better understand tick dispersal patterns using population genetic analyses. We analysed more than 1,000 ticks sampled at 32 sites in two conservation areas in Mozambique. Tick population sizes were relatively small, with signs of bottlenecks. Ticks from the two areas were completely isolated from each other, but within each area, ticks from nearby burrows were connected, probably because warthogs carry them from one burrow to another. These short-distance migration patterns shed light on virus dispersal and support the maintenance of buffer zones around protected areas.

---

**Keywords:** Vector surveillance, Soft tick, *Ornithodoros*, Population genetics, Microsatellite markers, African swine fever, Warthog

## Background

Vector-borne diseases have a major impact on human health, accounting for more than 17% of all human infectious diseases [1], but also pose a growing health risk to livestock as the size of livestock production systems increases [2]. A vector-borne disease is the result of complex interactions between the transmitted pathogen, the vector, the host and the environment. Therefore, population size, population dynamics, migration patterns and mating system of hematophagous arthropods represent critical information for understanding the epidemiology of vector-borne diseases and, consequently, for vector control programs and disease management strategies. However, much basic information about vector ecology is frequently unknown because their small size, location and behavior complicate direct observation of their population biology. Population genetics offers a way around these limitations, based on variation in allelic frequency at neutral genetic markers [3,4].

In veterinary medicine, ticks are the most important arthropod vector of pathogens responsible for animal diseases, ahead of mosquitoes [5]. Among these diseases, only a few are caused by viruses. This is the case with African swine fever (ASF), the most complex and devastating emerging infectious disease of pigs. In domestic pigs, infection can cause up to 100% mortality in about 10 days [6,7]. There is currently no approved treatment for this disease, while live attenuated vaccines have received regulatory approval in only a few countries, mainly in Southeast Asia. In the wild, the ASF virus has existed for millions of years in a sylvatic cycle involving warthogs (*Phacochoerus africanus*) and soft ticks (of the *Ornithodoros moubata* complex) that infest their burrows [8]. After infection, wild suids show no clinical signs, despite significant viremia [9,10]. The sylvatic cycle involving soft ticks represents a constant risk of ASF virus spillover to domestic pigs, particularly in the buffer zone of wildlife conservation areas. The link between the sylvatic and the domestic cycles is still poorly understood. Experimental evidence suggests that direct contact transmission between domestic pigs and warthogs is unlikely [9]. Therefore, either indirect contamination occurs with domestic pigs being potentially infected by warthog fomites, or soft ticks are involved in viral transmission from wild suids to pigs. Although there is no direct evidence of tick movement between warthog burrows and pigpens, some studies have reported the presence of soft ticks in pigpens near conservation areas [11]. Investigating dispersal patterns of soft ticks in Southern Africa is critical in order to assess their role in ASF virus transmission at the wild-domestic interface.

The *Ornithodoros moubata* complex consists of nine species of soft ticks distributed throughout Eastern and Southern Africa [12]. The presence of these ticks appears to be influenced by several factors, including soil properties, altitude and vegetation [8,12–14]. Among these species, *Ornithodoros phacochoerus* is found in the Eastern part of South Africa, in Swaziland and in Mozambique. In the latter country, the existence of both sylvatic and domestic cycles has been demonstrated [11,15], and ASF virus represents a high risk for pig farming systems [16].

To date, population genetics studies have predominantly focused on hard ticks [17] and the few related to soft ticks are based on mitochondrial markers [18–21], although specific markers have been developed for the soft tick *Ornithodoros coriaceus* [22]. Population genetics studies have shown that gene flow among tick populations is often strongly influenced by host movements but also by tick behaviour and life-history traits [17]. This is particularly relevant for nidicolous ticks, which remain associated with discrete nesting or resting sites and may therefore experience more spatially restricted gene flow than would be expected from host mobility alone. For example, population genetic studies of the nidicolous hard ticks *Ixodes uriae* and *Ixodes arboricola* have shown that genetic structure varies according to host species, nesting behaviour, and the spatial distribution of nesting sites [23,24]. A similar mechanism may operate in *O. phacochoerus*, which is closely associated with its hosts’ resting sites. However, unlike hard ticks, soft ticks take very short blood meals (less than 2 hours) and are therefore less likely to be dispersed by their host. Even if the presence of soft ticks has been reported on warthogs outside their burrows [25], it is not known whether this leads to efficient dispersal.

Using a set of 15 microsatellite markers selected from a panel of 19 loci previously developed from genomic data of *Ornithodoros porcinus* and *Ornithodoros moubata* [26], we studied the population genetic structure of *Ornithodoros phacochoerus* in Mozambique. With 32 sites sampled between two wildlife conservation areas, our objective was to evaluate the dispersal pattern of this soft tick to investigate its potential role in ASF virus spillover from the sylvatic cycle to the domestic cycle. This is the first population genetic study based on nuclear markers for soft ticks.

## Results

### Genetic diversity

Genetic diversity indices were computed for each of the 32 sampling sites (plus the 2 temporal replicates) across the two study areas, Coutada 9 Game Reserve (n = 22 sites + 1 replicate) and Gorongosa National Park (n = 10 sites + 1 replicate) (Table S1). Gorongosa sites consistently exhibited higher genetic diversity than Coutada 9 sites, with mean expected heterozygosity (H_E_) ranging from 0.35 to 0.526 in Gorongosa compared to 0.039 to 0.397 in Coutada 9. The mean number of alleles per locus followed the same pattern, ranging from 2.625 to 3.563 in Gorongosa versus 1.313 to 2.375 in Coutada 9. The high degree of allelic differentiation further supported this pattern, with 50 of the 82 alleles detected (61%) being private to one of the two conservation areas (15 in Coutada 9 and 35 in Gorongosa). Inbreeding coefficients (F_IS_) were predominantly positive across both conservation areas, indicating a widespread deficit in heterozygotes, but differed substantially in magnitude and dispersion between the two reserves. In Gorongosa, F_IS_ values were low and remarkably homogeneous, mostly ranging from 0.053 to 0.094 (with the exception of G6_24 at 0.227 and G3_24 at 0.000). In contrast, Coutada 9 exhibited both higher and more variable F_IS_ values (ranging from -0.433 to 0.342), with over a quarter of the sampled sites displaying marked heterozygote deficiencies exceeding 0.20. Hardy–Weinberg equilibrium was rejected (p<0.05) in several sites, with departures being markedly more prevalent in Gorongosa National Park (n=9 sites) than in Coutada 9 (n=4 sites).

### Relevant level of population structure

At the global scale, combining all 32 sites from both Coutada 9 Game Reserve and Gorongosa National Park, the DAPC scatter plot revealed a clear separation between the two conservation areas along the first discriminant axis, with all Coutada 9 sites forming a distinct cluster on one side and all Gorongosa sites on the other (Figure 1). This differentiation was strongly supported by the membership probability plot at K = 2 (see Figure S1 for K = 3 to 5), in which all individuals were assigned with probability 1.0 to their conservation area, with no admixture detected between the two areas.

**FIGURE 1:**
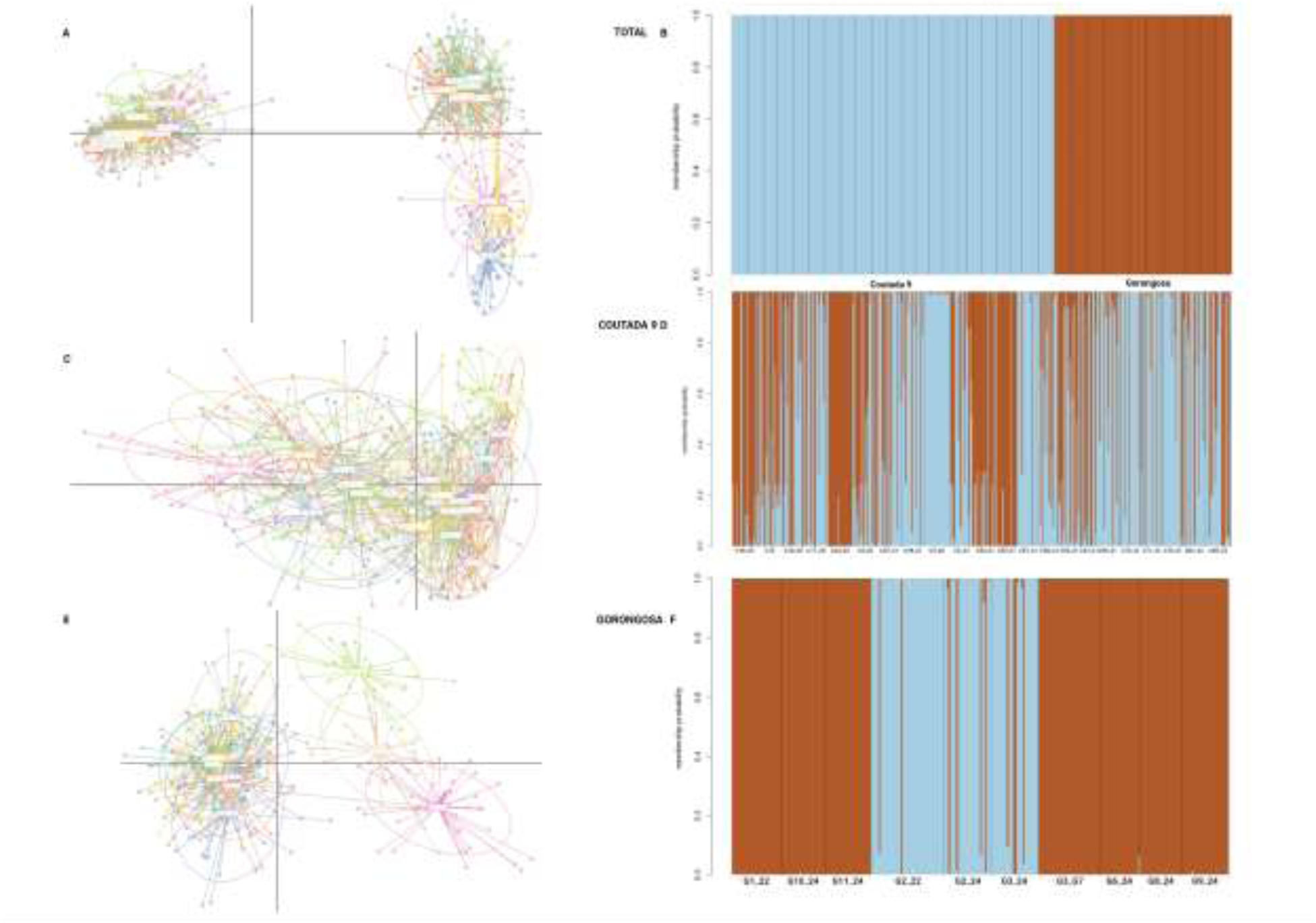
Sample distribution according to genetic proximity. A, C & E, Spatial discriminant analysis of principal components (sDAPC) scatterplot depicting the genetic structuring of individuals sampled across the different groups/sites. Each site is represented by one colour. B, D & F, DAPC plots for K = 2 subpopulations selected using *find.clusters* with 50 principal components retained. Each color represents one of the 2 subpopulations defined and the probability for each tick to belong to this subpopulation. Ticks are grouped by site with the site number under each group. A and B, analysis performed with all 32 sites. C and D, analysis performed with the 22 sites from Coutada 9 Game Reserve. E and F, analysis performed with the 10 sites from Gorongosa National Park.

Given the clear genetic differentiation observed between Coutada 9 Game Reserve and Gorongosa National Park at the global scale, all subsequent analyses were conducted separately for each protected area. Within the Coutada 9 Game Reserve, the DAPC scatter plot showed extensive overlap among all sampling sites, with no clear separation between groups. The membership probability plot at K = 2 displayed a high level of admixture across all sites, with most individuals exhibiting mixed assignment probabilities between the two inferred clusters, and no site showing consistent assignment to a single cluster. Within Gorongosa National Park, a more differentiated pattern was observed. The DAPC scatter plot revealed two partially distinct groupings, with sites G2_22, G2_24, and G3_24 separated from the remaining sites along the first discriminant axis. This spatial structuring was reflected in the membership probability plot at K = 2, where these sites were predominantly assigned to one cluster while the majority of remaining sites were largely assigned to the other, with only limited admixture between the two genetic groups.

### AMOVA

Conservation area was the primary driver of genetic structure, explaining 51.2% of total variation (Φ-PT = 0.512), with an additional 8.0% attributed to differentiation among populations within areas. Temporal variation among sampling years also accounted for a substantial proportion of genetic differentiation (29.1%, ΦCT = 0.291, with 19.0% among populations within years). In contrast, habitat type explained a negligible proportion of between-group variance (1.9%, ΦCT = 0.019) (Table S2).

### Isolation by distance

Pairwise fixation index (F_ST_) estimates revealed marked genetic structure across the sampled populations (Figure 2). Pairwise F_ST_ values between Coutada 9 and Gorongosa conservation areas were consistently high, ranging from 0.354 to 0.766 with a mean inter-parc F_ST_ of 0.554. In contrast, within-areas differentiation was substantially lower averaging 0.174 within Coutada 9 and 0.151 within Gorongosa. This pattern was fully corroborated by Jost’s actual differentiation, which exhibited pronounced divergence between the two conservation areas (mean: 0.651), compared to minimal differentiation within Coutada 9 (0.065) and Gorongosa (0.136) (Figure S2).

**Figure 2:**
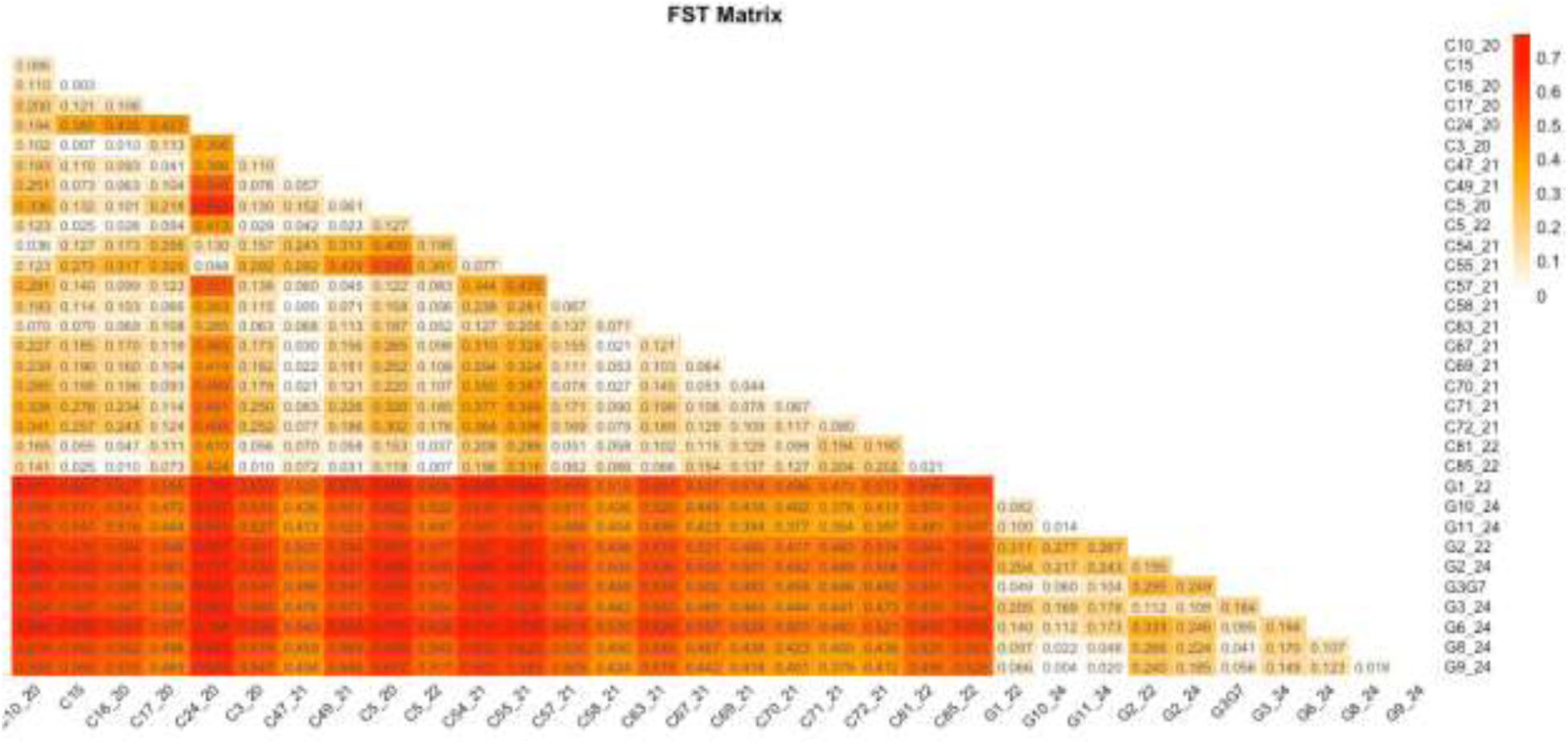
Pairwise Weir and Cockerham’s F_ST_ values are indicated within each cell and visualised along a colour gradient from white (F_ST_=0, no differentiation) to dark red (F_ST_>0.7, strong genetic differentiation). Labels with the prefix “C” correspond to sampling localities in Coutada 9, and “G” to localities in Gorongosa National Park.

In Coutada 9, significant isolation-by-distance was detected among sites (b=0.131, Mantel test, 9,999 permutations, r=0.101, p<0.0001) (Figure S3). In Gorongosa National Park, an isolation-by- distance pattern was also detected (b=0.115, Mantel test, 9,999 permutations, r=0.253, p=0.0005).

### Effective population size

Effective population size (N_e_) calculation using NEESTIMATOR with the co-ancestry method ended up with 1/34 infinite values whereas the linkage disequilibrium method yielded 4/34 infinite values for Coutada 9 and 2/34 for Gorongosa (Table S3). For the temporal method of calculation, two years between each sampling were used, corresponding to two generations of ticks. For Coutada 9 (site 5_20/5_22), the three methods yielded N_e_ estimates of 3.8 (Pollak), 4.3 (Nei/Tajima), and 6.3 individuals (Jorde/Ryman), giving a mean N_e_ of 4.8. For Gorongosa (site G3_22/G7_24), the estimates were 2.7 (Pollak), 3.1 (Nei/Tajima), and 2.5 individuals (Jorde/Ryman), giving a mean N_e_ of 2.8.

Using COLONY (sibship reconstruction), N_e_ estimates in Coutada 9 ranged from 5 to 351, with a median of 15.0 and an unweighted mean of 29.5 (driven upwards by a single outlier, population C24_20, with N_e_=351 and an upper 95% confidence interval reaching infinity) (Figure 3). In Gorongosa, N_e_ estimates showed higher and more consistent values, ranging from 11 to 36, with a median of 21.0 and an unweighted mean of 21.8. The test of moderators revealed a significant difference in effective population size between the two groups (QM=7.84, df=1, p=0.0051). The model accounted for a large proportion of structured between-group heterogeneity (R²=79.48%, τ2=0.0150, I2=10.24%). The summary effective population size was estimated at N_e_=14.9 [12.6;17.6] for Coutada 9, compared to N_e_=21.7 [17.0;27.6] for Gorongosa.

**Figure 3:**
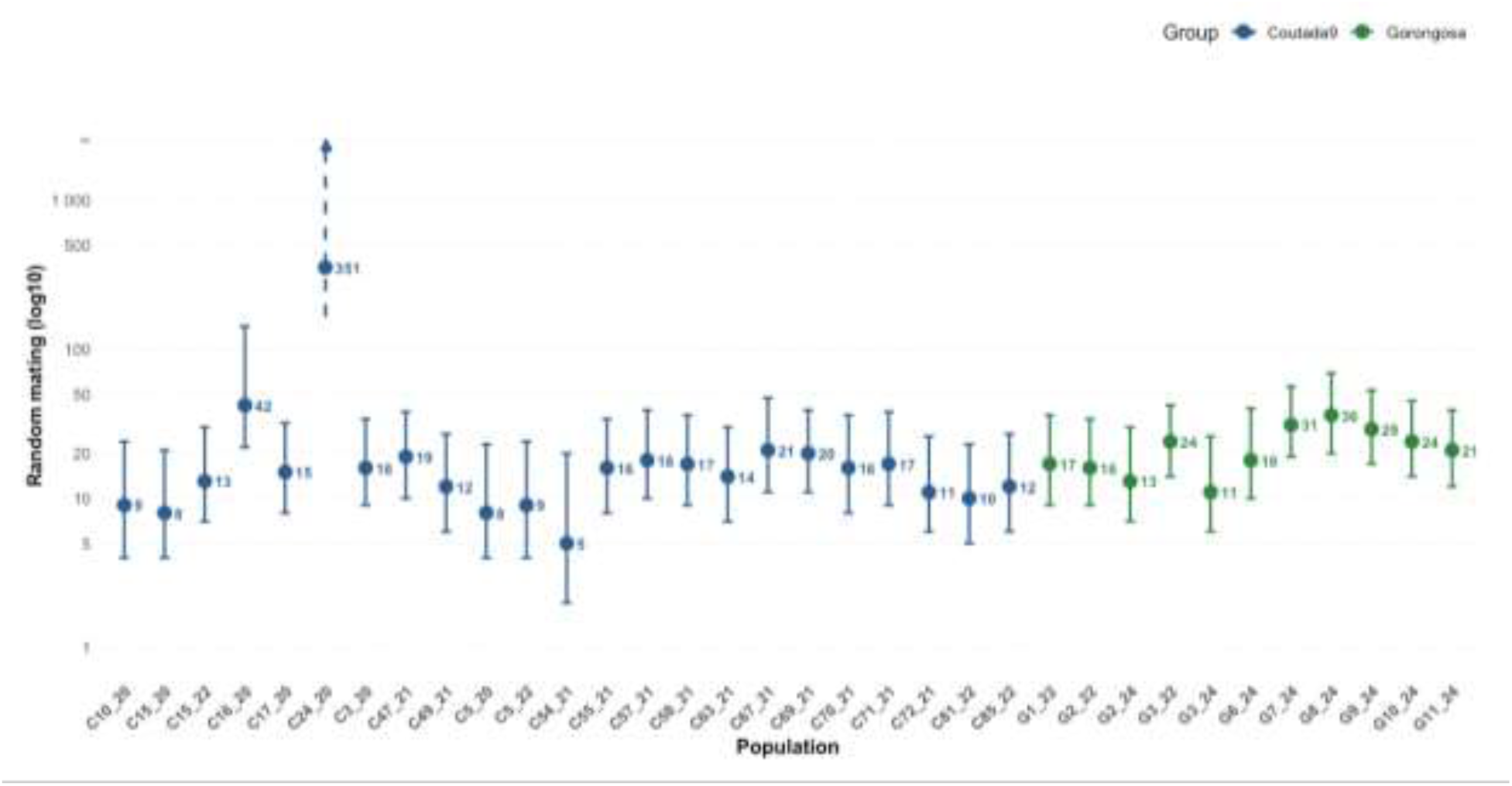
Effective population size estimates across sampled populations of *Ornithodoros phacochoerus*. Estimates were inferred using the sibship assignment method in COLONY under a random mating model with polygamy and inbreeding allowed. Filled circles indicate point estimates (N_e_, with values displayed next to each point), and vertical error bars represent the corresponding 95% profile likelihood confidence intervals on a log10 scale. Colors distinguish populations sampled in Coutada 9 (blue) and Gorongosa National Park (green). The dashed arrow for population C24_20 denotes an upper 95% confidence interval bound reaching infinity (∞).

In Coutada 9, neighbourhood size was estimated at 7.6 individuals and the number of migrants per generation at 1.21 individuals. Based on the meta-analytic estimate of N_e_=14.9, the migration rate was estimated at m=0.08. In Gorongosa, neighbourhood size was estimated at 8.63 individuals and the number of migrants per generation at 1.37 individuals. Based on the meta-analytic estimate of N_e_=21.7, the migration rate was estimated at m=0.06. Using a density of burrows of 16 sites/km², the average dispersal distance per generation was 0.1 km in Coutada 9 and 0.08 km in Gorongosa.

### Bottleneck

Per-site Wilcoxon sign-rank tests were conducted under three mutation models (IAM, TPM, and SMM) to detect recent demographic bottlenecks in populations from Coutada 9 Game Reserve and Gorongosa National Park (Table S4). In Coutada 9, 10 out of 23 sites showed significant p- values under IAM, dropping to four under TPM and three under SMM. In Gorongosa, bottleneck signals were stronger and more consistent across mutation models. Eight out of 11 sites were significant under IAM, six under TPM and two under SMM.

## Discussion

### Two conservation areas genetically separated and differentiated

This study is the first to investigate the population structure of a tick vector of the African swine fever virus in an endemic area. The first striking genetic characteristic revealed by our study is the very marked differentiation between the two conservation areas, as previously suggested [26]. The mean pairwise F_ST_ between Gorongosa and Coutada 9 sites is 0.55, indicating strong genetic differentiation and restricted gene flow between the two areas. This high F_ST_ value should be interpreted with caution since it can be largely influenced by the low level of genetic diversity in Coutada 9. However, we also found high Jost’s differentiation between Gorongosa and Coutada 9, a measure that is less dependent on within-population diversity than F_ST_ [27]. The concordance between the two measures provides complementary evidence for strong genetic differentiation between the two conservation areas, rather than being solely attributable to differences in within-area genetic diversity. The high degree of allelic differentiation further supports this pattern, with 50 of the 82 alleles detected being private to one of the two conservation areas. Such a high proportion of private alleles is consistent with limited genetic exchange between the two areas and may reflect the combined effects of restricted gene flow and genetic drift.

This genetic differentiation is consistent with the marked geographic separation between Gorongosa National Park and Coutada 9 Game Reserve, which are approximately 150 km apart in a straight line between sampled sites. While the warthog populations are relatively well characterized in the two conservation areas, there is currently no documented evidence of warthog presence between the two areas, possibly as a result of poaching pressure outside conservation areas and human settlements. The Macossa-Tambara Ecosystem, established in 2024, brings together the neighbouring Coutada 9 and Coutada 13 concessions, with the latter lying between Coutada 9 and Gorongosa, with the aim of restoring and connecting natural habitats. Such conservation efforts may contribute to increased wildlife connectivity and, potentially, to future tick gene flow at a larger spatial scale.

Genetic clustering identified two genetic clusters corresponding to two separated geographic areas in Gorongosa whereas all individuals in Coutada 9 were assigned to a single cluster. This difference in genetic structure between the two areas may partly be related to the lower genetic diversity observed in Coutada 9 (despite a larger number of sites sampled). Indeed, reduced allelic variation may limit the power of clustering methods to detect fine-scale genetic structure. However, we obtained moderate levels of genetic differentiation among Coutada 9 sites, indicating that genetic structure is present but does not translate into distinct clusters. While Coutada 9 is mostly characterized by wooded savanna (miombo), Gorongosa encompasses a more heterogeneous savanna-woodland-floodplain landscape. Even if the genetic clusters in Gorongosa are not clearly linked with landscape variation, this greater environmental heterogeneity may have contributed to the emergence or maintenance of genetic structure.

### A metapopulation system

Within each conservation area, tick populations were strongly structured by sampling site, with significant genetic differentiation among sites and a clear signal of isolation-by-distance. Genetic differentiation increased with geographical distance, suggesting that gene flow may occur predominantly among neighbouring sites, potentially following a stepping-stone pattern across patchily distributed habitats. This spatially structured gene flow may contribute to the formation of differentiated genetic clusters, as observed in Gorogonsa. Such isolation-by-distance and stepping-stone pattern has already been observed in the endophilic hard tick *Ixodes uriae* associated with seabirds [23].

Estimates of effective population size, N_e_, varied substantially depending on the estimation method, sampling sites and conservation areas. Overall, the sibship method implemented in COLONY yielded lower N_e_ estimates than the linkage disequilibrium method. In addition, the linkage disequilibrium method returned infinite point estimates or upper confidence limits for many populations. An infinite estimate may reflect either a very large N_e_ or, more likely, insufficient statistical power (limited sampling size or low number of molecular markers) as no linkage disequilibrium signal above that expected from sampling error could be detected.

Differences in N_e_ estimates among methods have previously been reported (e.g. Lévêque et al. 2024). Based on the sibship method, the summary N_e_ were between 10 and 30 ticks per site. To our knowledge, this is the first report of effective population size in soft ticks, and estimates of N_e_ remain scarce even among hard ticks. Using the linkage disequilibrium method and temporal methods, Huber et al. (2019) reported similarly low estimates for *Amblyomma variegatum*, a three-host tick found primarily on cattle, whereas larger estimates have been reported for *Rhipicephalus microplus* and *Ixodes ricinus* [30,31].

Estimation of N_e_ relies on several simplifying assumptions, including limited or absent migration among populations. Substantial gene flow among sampling sites can therefore violate these assumptions and potentially lead to upwardly biased estimates of N_e_, as migration increases the number of potential parents contributing to the sampled population [32]. Despite these methodological limitations, the consistently low N_e_ estimates obtained with the sibship method suggest that effective population sizes of *Ornithodoros phacochoerus* may be relatively small. This interpretation is also consistent with the low number of adults collected at each site compared with the number of nymphs.

Evidence of recent bottlenecks was detected in several populations, with approximately 20% of sites within each conservation area displaying a bottleneck signature under all three models tested. Combined with the observed isolation-by-distance pattern, relatively small effective population sizes, and high migration rates among sites, these results are consistent with a metapopulation structure characterized by local population turnover and potentially recurrent extinction–recolonization dynamics. In such systems, the persistence of individual patches may depend on demographic and environmental stochasticity. In Gorongosa, many sites are subjected to strong seasonal flooding, while in Coutada 9 bushfires constitute an important disturbance. These phenomena could contribute to extinction of tick populations, but this would require dedicated work to confirm this hypothesis. This metapopulation model provides a framework for understanding the highly unpredictable occurrence of ticks among burrows and resting sites, including the approximately 30% of burrows in which no ticks were detected, as recently reported in Coutada 9 by Lameira et al. (2026).

Based on effective population size estimates (using COLONY) and the isolation-by-distance regressions, we further estimated migration rates and average dispersal distance per generation (about 7% and 100 m). These estimates suggest that gene exchange among *O. phacochoerus* populations occurs at a relatively local spatial scale, with non-negligible connectivity among neighbouring populations. Short distance connectivity has already been demonstrated in *Rhipicephalus microplus* in New-Caledonia, where tick population displayed a pattern of isolation-by-distance with migration distance estimated at a few hundred meters per tick generation, presumably driven by within-herd cattle rotation among pastures [31]. Many genetic studies of ectoparasites have highlighted the influence of host movements on parasite dispersal and population structure [24,33,34] with even higher dispersal in parasites than in their hosts (reviewed in Mazé-Guilmo et al. 2015). In the present system, the density and movement of hosts likely plays an important role in shaping soft tick population structure, despite very short blood meal duration on the host compared to hard ticks.

### Tick populations structured by their hosts

Soft ticks of the *O. moubata* complex can feed on a wide variety of hosts. Collection of specimens on animals as well as blood meal analyses identified warthog, antbear and porcupine as primary hosts and tortoise hyena, jackal, pangolin as secondary hosts [13,14,36]. Those ticks are also associated with human and domestic animals (dogs, fowls). In this study, ticks were collected mostly in burrows used by warthogs, aardvarks, and porcupines. The spatial distribution of occupied burrows may facilitate gene flow among tick populations. Based on published data, burrow density in southern African savannas appears to be on the order of 15–55 burrows per km², although the proportion of burrows occupied is substantially lower and varies temporally [37–39]. A density of approximately 16 occupied burrows per km² would correspond to an average distance of approximately 250 m between burrows, assuming a regular spatial distribution. In addition, home ranges of warthogs, aardvarks and porcupines are similarly restricted, ranging from 0.64 to 3.41 km², 1.3 to 3.5 km² and 0.4 to 3.5 km², respectively [37,40–43]. These species can have overlapping home ranges and individuals use multiple burrows within their home range. The combination of these spatial scales and movement patterns suggests that neighbouring burrows may be frequently connected by host movements, potentially facilitating local tick dispersal. Thus, *Ornithodoros phacochoerus* may have opportunities for dispersion between nearby burrows, which is consistent with the genetic structure described here and with the rough estimate of dispersal distance per generation (approximately 100 meters). In addition, warthogs, aardvarks and porcupines switch between burrows over time [38,40,41], and this translates into a large proportion of burrows remaining temporarily unoccupied by their host. Such temporal heterogeneity in burrow occupancy could contribute to the local extinction as suggested by the bottleneck signals in several tick populations.

Bottleneck events may also have occurred at broader spatial and temporal scales in response to substantial fluctuations in host population sizes. In Mozambique, many conservation areas experienced severe wildlife depletion during the second half of the 20th century, particularly during the colonial period and the subsequent civil war [44–46]. In Gorongosa National Park, populations of most large mammal species declined by more than 90% during the war period [47]. Following the end of the civil war, increased protection and conservation efforts allowed many large-mammal populations to recover. In particular, the estimated density of warthogs based on aerial surveys increased from approximately 0.3 individuals per km² in 1997 to 6.2 individuals per km² in 2018. In Coutada 9 Game Reserve, a comparable increase in warthog abundance was reported, from 338 individuals counted in 2005 to 3500 in 2024 (https://www.patrolling.org/coutada-9-mozambique/). No warthog reintroductions were conducted in either conservation area during this period, suggesting that these increases primarily reflect natural population recovery.

Such large fluctuations in host abundance may have also affected their associated parasite populations. However, bottleneck tests are designed to be applied to individual populations assumed to be panmictic and therefore cannot be directly applied to a metapopulation or to an assemblage of genetically differentiated populations such as those occurring within a conservation area. We can therefore only hypothesize that the low genetic diversity observed in Coutada 9 may partly reflect a historical loss of genetic diversity following a major demographic contraction in *O. phacochoerus* populations associated with the severe decline in host abundance during the period of conflict.

### Application to African Swine Fever Virus control

Like other species of the *O. moubata* complex, *Ornithodoros phacochoerus* is a vector for the African swine fever virus [8], and both Coutada 9 Game Reserve [48] and Gorongosa National Park [15] have been confirmed to have a sylvatic cycle for African swine fever virus. Because of the tick migration associated with the frequent movements of warthogs between burrows, it is highly likely that ticks are moving ASF virus from burrows to burrows and from warthog group to another, thus being an active actor in the ASF sylvatic cycle. Furthermore, soft ticks should be able to transmit the virus from the sylvatic cycle to the domestic cycle. Since we were not able to find ticks in pigpens around Coutada 9 Game Reserve, it was not possible to properly tackle this issue. Nonetheless, based on the estimated average dispersal distance per generation (about 100 meters) calculated in this study and estimates of host home ranges, the viral transmission to pigs by soft ticks would occur if free-ranging pigs or unfenced pigpens were present in close proximity to the conservation areas. This has already been suspected in Gorongosa National Park, where pigs near the park were infected with the same ASF genotype as ticks and warthogs from inside the park [15]. The short-estimated migration distance per generation provides a quantitative indication of the spatial scale over which local tick connectivity may occur and could complement the larger-scale zoning and biosecurity measures currently used in ASF-endemic areas of southern Africa, which primarily aim to prevent contact between domestic pigs, wild suids and soft ticks [49].

## Methods

### Tick sampling

*Ornithodoros phacochoerus* ticks were collected in Mozambique from two different wildlife conservation areas: the Coutada 9 Game Reserve (4 333 km²) from the district of Macossa, Manica province, and the Gorongosa National Park (4 067 km² for the core protected area, 5 333 km² for the buffer zone) from the district of Gorongosa, Sofala province (Figure 4). Ticks were collected over four sampling campaigns: in October 2020, 2021 and 2022 for Coutada 9 and in October 2022 and October 2024 for Gorongosa (Table 1). Ticks from Coutada 9 Game Reserve were collected from 22 different sites (including one site, sampled in 2020 and voluntarily resampled in 2022, identified as C5_20 and C5_22 respectively). Pigpens (n=11) located in the buffer zone of the Coutada 9 Game Reserve were also investigated but no ticks were collected in these sites. Ticks from Gorongosa National Park were collected from ten different sites (including one site, sampled under a cabin deck in 2022 and voluntarily resampled in 2024, identified as G3_22 and G7_24 respectively).

**Figure 4:**
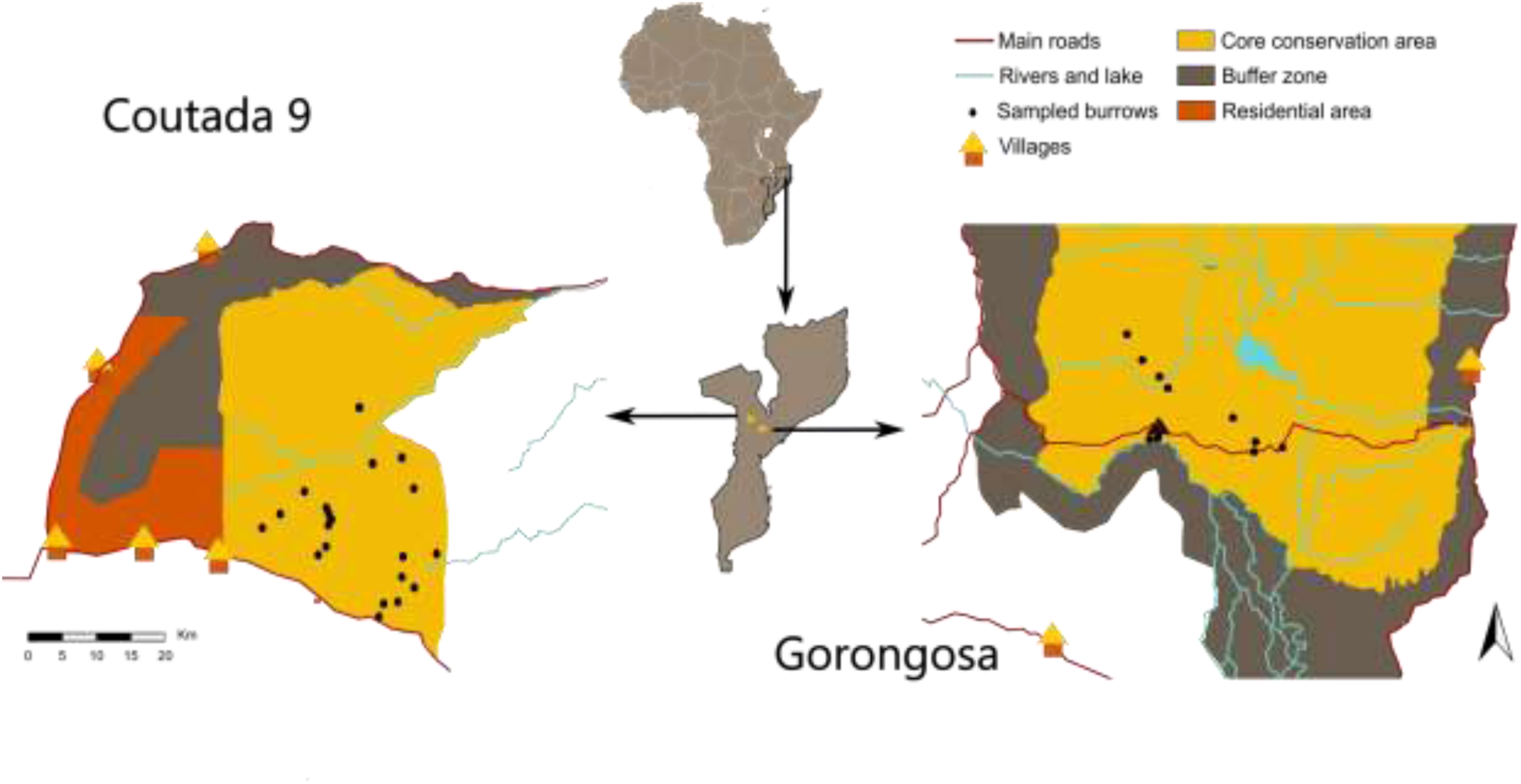
Location of sampling sites (black spots) for *Ornithodoros phacochoerus* in Coutada 9 Game Reserve and Gorongosa National Park, Mozambique. Coutada 9 Game Reserve map was based on a map provided by Mokore Wildlife. Gorongosa National Park map was derived from https://www.biointeractive.org/classroom-resources/gorongosa-national-park-interactive-map.

**Table 1:** Sites collected within each conservation area. For each site, we indicate the type of site, gps location, date of sampling (DD/MM/YYYY), number of ticks sampled and number of ticks retained for analyses (genotype with less than 4 missing loci). Some sites share the same GPS location data because the burrows occur in close proximity to each other. Only C5_20 / C5_22 in Coutada 9 and G3_22 / G7_24 are temporal replicates of the same site.

| Site ID | Conservation area | Habitat | Latitude | Longitude | Date | Nber of ticks sampled | Nber of ticks analyzed |
|---|---|---|---|---|---|---|---|
| C3_20 | Coutada 9 | Rocks | -17.65284 | 33.74459 | 29/09/2020 | 30 | 28 |
| C5_20 | Coutada 9 | Rocks | -17.66802 | 33.74428 | 29/09/2020 | 30 | 30 |
| C10_20 | Coutada 9 | Rocks | -17.62435 | 33.71297 | 30/09/2020 | 30 | 30 |
| C15_20 | Coutada 9 | Rocks | -17.6609 | 33.74788 | 31/09/2020 | 30 | 29 |
| C16_20 | Coutada 9 | Rocks | -17.6609 | 33.74788 | 31/09/2020 | 30 | 22 |
| C17_20 | Coutada 9 | Burrow | -17.6941 | 33.74793 | 31/09/2020 | 30 | 30 |
| C24_20 | Coutada 9 | Burrow | -17.67184 | 33.65805 | 03/10/2020 | 30 | 27 |
| C49_21 | Coutada 9 | Rocks | -17.64567 | 33.7415 | 12/10/2021 | 30 | 26 |
| C54_21 | Coutada 9 | Burrow | -17.6538 | 33.68138 | 12/10/2021 | 30 | 27 |
| C55_21 | Coutada 9 | Burrow | -17.51519444 | 33.78458333 | 12/10/2021 | 30 | 23 |
| C57_21 | Coutada 9 | Burrow | -17.58047222 | 33.84 | 12/10/2021 | 30 | 24 |
| C58_21 | Coutada 9 | Burrow | -17.62055556 | 33.85561111 | 12/10/2021 | 30 | 24 |
| C63_21 | Coutada 9 | Burrow | -17.70902778 | 33.72416667 | 13/10/2021 | 30 | 26 |
| C67_21 | Coutada 9 | Burrow | -17.78730556 | 33.81016667 | 15/10/2021 | 30 | 16 |
| C71_21 | Coutada 9 | Burrow | -17.70919444 | 33.84152778 | 16/10/2021 | 30 | 25 |
| C47_21 | Coutada 9 | Burrow | -17.77041667 | 33.81638889 | 11/10/2021 | 30 | 29 |
| C69_21 | Coutada 9 | Burrow | -17.76808333 | 33.83475 | 15/10/2021 | 30 | 28 |
| C70_21 | Coutada 9 | Burrow | -17.74963889 | 33.85616667 | 15/10/2021 | 30 | 25 |
| C72_21 | Coutada 9 | Burrow | -17.70533333 | 33.88519444 | 16/10/2021 | 26 | 20 |
| C81_22 | Coutada 9 | Burrow | -17.588093 | 33.8019 | 25/10/2022 | 30 | 29 |
| C85_22 | Coutada 9 | Burrow | -17.660785 | 33.748075 | 24/10/2022 | 30 | 29 |
| C15_22 | Coutada 9 | Rocks | -17.6609 | 33.74788 | 25/10/2022 | 30 | 24 |
| C5_22 | Coutada 9 | Rocks | -17.66802 | 33.74428 | 25/10/2022 | 30 | 26 |
| G1_22 | Gorongosa | Burrow | -18.979355 | 34.345292 | 28/10/2022 | 45 | 31 |
| G2_22 | Gorongosa | Burrow | -19.009308 | 34.4881 | 28/10/2022 | 51 | 46 |
| G3_22 | Gorongosa | Deck | -18.977531 | 34.352182 | 28/10/2022 | 42 | 38 |
| G2_24 | Gorongosa | Burrow | -19.007218 | 34.524308 | 30/09/2024 | 30 | 30 |
| G3_24 | Gorongosa | Burrow | -19.011393 | 34.487877 | 30/09/2024 | 30 | 30 |
| G6_24 | Gorongosa | Burrow | -18.96342 | 34.455017 | 01/10/2024 | 30 | 26 |
| G7_24 | Gorongosa | Deck | -18.977531 | 34.35182 | 01/10/2024 | 30 | 29 |
| G8_24 | Gorongosa | Burrow | -18.919788 | 34.358687 | 02/10/2024 | 30 | 27 |
| G9_24 | Gorongosa | Burrow | -18.902673 | 34.346868 | 02/10/2024 | 30 | 29 |
| G10_24 | Gorongosa | Burrow | -18.878675 | 34.323774 | 04/10/2024 | 30 | 28 |
| G11_24 | Gorongosa | Burrow | -18.841433 | 34.299777 | 04/10/2024 | 30 | 29 |

There were two different types of sites: “rocks” were shady areas made of rock and mud, where wild herbivores, including warthogs, spent time resting, “burrows” were holes in the ground that were primarily dug by aardvarks (*Orycteropus afer*) and used by warthogs to sleep and rear their young, thus escaping predation (Table 1 and Figure S4). One exception in Gorongosa National Park was under a cabin in Chitengo camp which was collected after observation of warthogs using the site for resting (identified as G3_22 and G7_24, for the two temporal replicates).

The selection of sites for genotyping was based on the number of collected ticks, with a target sample size of 30 ticks per site. Tick samplings were performed by excavating the soil from the side and the entrance of the burrow or by collecting soil from resting areas, then by spreading this material over a black tarp under sunlight. Ticks were then collected by hand and kept in a tube with some sand until DNA extraction. Tick DNA was shared with the Agricultural Research Institute of Mozambique (IIAM) under Material Transfer Agreement and in compliance with the Nagoya Protocol.

### DNA extractions and molecular identification

Between 26 and 51 individuals were collected per site (mean=31.1, SD=5), and DNA was extracted from each individual (Table 1). Tick washing and DNA extraction were performed following the method previously described in Taraveau et al. (2024). Based on sequencing of 12S (Genbank QB168940-QB168941) and 16S (Genbank QB160004-QB160007) rRNAs, ticks were identified as *O. phacochoerus*.

### Microsatellite characterization and genotyping

Genotyping was performed by capillary electrophoresis at the GPTR Laboratory (Great Regional Technical Platform of Genotyping, AGAP Institut/CIRAD, Montpellier, France) with an ABI 3500xL Genetic Analyzer (Applied Biosystems, Foster City, CA, USA), using microsatellite markers already published with the same protocol of DNA amplification [26]. Alleles were scored using GENEMAPER® v.6 software (Applied Biosystems, Waltham, MA, USA). Allele scoring was performed automatically according to the bin set previously designed for the markers, then manually checked by two different experimenters.

We first genotyped 686 ticks from 22 sites in Coutada 9 and 138 ticks from 3 sites in Gorongosa using the 19 microsatellites already selected. To test the quality of markers, the frequency of null alleles was estimated using FreeNA, accounting for the proportion of non-amplifying individuals (putative null homozygotes) at each locus, with 10,000 bootstrap replicates [50]. Then, a one-sided Spearman’s correlation test was also performed in R version 4.2.3 (2023-03-15) [51] to evaluate if the residual positive F_IS_ (inbreeding coefficient) in each locus could be explained by the number of blanks used as a signal for null alleles (S=614.95, p-value=0.7245). Finally, MICROCHECKER [52] was used to evaluate the number of populations affected by the presence of null alleles, stuttering or short allele dominance in each locus. Based on the collected data, loci deviating for Hardy-Weinberg equilibrium (HWE) were discarded.

Three microsatellite markers (ms-61, ms-76, and ms-111) were discarded due to signal of null alleles or stuttering associated with positive F_IS_, and significant deviation from HWE (Table S5). No signature of short allele dominance was detected in any of the loci and they were considered independent. Ms-87 was not used in the subsequent genotyping, leading to a total of fifteen loci genotyped. In this new set of markers, the concentration of each primer within each multiplex was adjusted to improve genotyping (Table S6). Genotyping was performed on 7 new sites and one replicated site in Gorongosa, representing 232 additional individuals genotyped. A total of 1056 ticks were genotyped, 754 of which presented full genotypes for all loci and 293 partial genotypes. Individuals with more than 4 missing loci were excluded from the analysis, leading to a total of 940 individuals with a mean of 27.65 individuals per site (range= [16;46], SD=4.95, Table 1).

Analyses were performed using the full dataset including temporal replicates except for genetic structure analyses (DAPC, AMOVA, isolation-by-distance) for which we kept only the first temporal replicate for sites sampled twice.

### Genetic diversity and differentiation

Genetic diversity was assessed for each population using the R package *hierfstat* [53]. For each population, we calculated the observed heterozygosity (H_O_), the expected heterozygosity (H_E_) following Nei’s (1978) unbiased estimator, and the mean number of alleles per locus. The inbreeding coefficient (F_IS_) was estimated following Weir and Cockerham (1984), and 95% confidence intervals were obtained by bootstrap resampling over loci (1,000 permutations) using the *boot.ppfis* function. Departures from HWE were tested for each population across all loci using exact tests implemented in the *pegas* package [56], and a combined p-value was obtained using Fisher’s method. For each population, a one-sided Wilcoxon signed-rank test was applied across loci to assess whether H_O_ was significantly lower than H_E_, which would be consistent with a Wahlund effect resulting from cryptic population subdivision. Private alleles specific to each conservation area were identified using the *private_alleles* function from the *poppr* package [57].

Population genetic structure was investigated using Discriminant Analysis of Principal Components (DAPC), implemented in the *adegenet* package [58,59]. A first DAPC was conducted using predefined sampling localities as prior group membership. To identify genetic clusters independently of sampling localities, the *find.clusters* function was used to apply k-means clustering on the principal components of the genotypic data, exploring K = 2-23 clusters with 50 PCs retained prior to clustering. The optimal number of clusters was evaluated based on mean and minimum inter-cluster distances in the discriminant space, individual assignment success rates, and the number of statistically significant discriminant axes assessed by one-way ANOVA. As a complementary approach, k-means clustering was also applied on principal components derived from a PCA of the scaled genotypic matrix using *dudi.pca* [60]. Membership probabilities for each value of K were visualised as stacked bar plots with individuals ordered by sampling population.

To partition genetic variance among hierarchical levels, analyses of molecular variance AMOVA [61] were performed using the *poppr.amova* function from the *poppr* package. Three hierarchical groupings were tested successively: (i) conservation area, ii) year of sampling and (iii) type of sampling site.

Pairwise genetic differentiation among populations was estimated using Weir and Cockerham’s (1984) F_ST_, computed with the *pairwise. WCfst* function in *hierfstat*. Confidence intervals were obtained by bootstrap resampling over loci (5,000 permutations), and pairwise F_ST_ values whose 95% CI overlapped zero were considered non-significant. In addition, we calculated Jost’s D_est_ measure of differentiation [62]. This statistic is based on variance in allele frequencies among populations and is independent of within population diversity. Jost’s D_est_ was computed using the *mmod* package.

Isolation-by-distance was tested using a Mantel test (9,999 permutations) between the matrix of pairwise F_ST_ and the matrix of geographic distances, as implemented in the *vegan* package [63]. Geographic distances between populations were calculated from mean sampling coordinates using the Haversine formula using the *geosphere* package [64]. Following Rousset (1997), linearised F_ST_ values (F_ST_/(1−F_ST_)) were regressed against the natural logarithm of geographic distance (km) to estimate the slope of the isolation-by-distance relationship (b). Under a two-dimensional model, the neighbourhood size is estimated as 1/b and the migration rate is 1/(2πbN_e_) where N_e_, the effective population size per site, is estimated independently (see below) [65,66]. A rough proxy of the average dispersal distance per generation was computed as 2√(1/(4πbN_e_D_sites_)) where D_sites_ is the density of sites (per km²) [67]. Field observations of the number of sites (burrows) effectively used by warthogs suggest densities of 15 sites/km² in the Sengwa Wildlife Research Area (Zimbabwe) [37] and 17 sites/km² in Hluhluwe-Umfolozi Park (South-Africa) [38]. Assuming that these estimates are applicable to Mozambique, we used a mean value of D_sites_=16 sites/km².

### Effective population size and bottleneck

Effective population size was calculated using NEESTIMATOR version 2.1 [68] and COLONY v.2.0.7.2 [69]. NEESTIMATOR was used with two different methods: one using all populations, and the other using only the two sites with two temporal replicates (C5_20 / C5_22 sampled in 2020 and 2022 in Coutada 9 and G3_22 / G7_24, sampled in 2022 and 2024 in Gorongosa). First, N_e_ per population was estimated using the co-ancestry model, with Jackknife for confidence intervals [70]. Second, N_e_ was also estimated using the linkage disequilibrium method for random mating with a lowest allele frequency value of 0.050 [71]. Finally, the temporal method was used with Pollak [72], Nei/Tajima [73], and Jorde/Ryman [74] methods, on the site number 5 which was sampled twice in Coutada 9 [75]. Two years separated the two sampling, which was evaluated as two generations of ticks between each sampling [76].

COLONY uses a maximum likelihood sibship reconstruction method [69] to estimate effective population size for each sampled population. We implemented the full-likelihood analysis method dedicated to dioecious diploid species with medium-length runs and high-precision likelihood calculations. The mating system was configured assuming polygamy for both males and females and no sibship prior, with random mating and inbreeding allowed. Inbreeding was enabled to account for the nidicolous lifestyle of *O. phacochoerus*. For each population, point estimates of N_e_ along with their associated 95% confidence were retrieved and log-transformed to stabilize variance and normalize distributions. To explicitly account for estimation uncertainty, we fitted an inverse-variance weighted mixed-effects meta-regression model using the *metafor* package in R [77]. Between-population residual heterogeneity (τ2) was estimated using the Restricted Maximum Likelihood estimator. Statistical significance of the conservation area effect was assessed via the Wald-type test of moderators (QM).

Signature of a bottleneck effect was investigated using the BOTTLENECK software version 1.2.02 [78] using the infinite allele model (IAM), the stepwise mutation model (SMM) and the two-phase mutation model (TPM). The TPM was set with default values (70% SMM, 30% IAM and variance of 30). To determine whether populations exhibited a significant excess of heterozygosity, p-values were calculated using a one-tailed Wilcoxon signed-rank test.

## Author contributions

FT coordinated the study, collected the samples in the field, performed the experiments, analyzed the data, and wrote the manuscript. CG analyzed the data, and wrote the manuscript. DB performed the experiments, and analyzed the data. CQ obtained the funding, coordinated the study, provided the material, and collected the samples in the field. TP obtained the funding, and coordinated the study. MJ, MD, and EL collected the samples in the field, and performed the experiments. AA performed the experiments. AF and JC collected the samples in the field. KH wrote the manuscript. HJ-P obtained the funding, conceived and coordinated the study, provided the material, collected the samples in the field, performed the experiments and wrote the manuscript. All authors read and corrected the final manuscript.

## Acknowledgements

The authors would like to thank the students from the IIAM who helped for sample sorting and DNA extractions, Edna Manuel Meque, Isaque António Francisco, Matias José Dina, Neidy Alfredo Lopes, Albertina Jaime Chabissa and Luisa Duarte Madeira, as well as Estelle Pineau who helped for tick sampling and DNA extractions. We would like to thank Mokore and Western camps which provided help and shelter during sampling in Coutada 9 Game Reserve. We would like to thank Gorongosa National Park for permission and help to conduct research (scientific permit PNG/DSCI/C235/2022). We also would like to thank Ronan Rivallan for the help on the GPTR platform, Thierry De Meeûs for great advice in the genotyping analysis and Alban Faure who read and corrected the first version of this manuscript.

## Data, scripts and codes availability

Data supporting the findings presented here are available within this paper and its supplementary material, genotyping dataset for the populations studied here will be available on Dryad repository shortly after publication.

