## Supplementary figures and tables S1 to S6 for "Isolation by distance and metapopulation structure in Mozambican populations of the soft tick *Ornithodoros phacochoerus*, a vector of African Swine Fever virus"

### Supporting Information

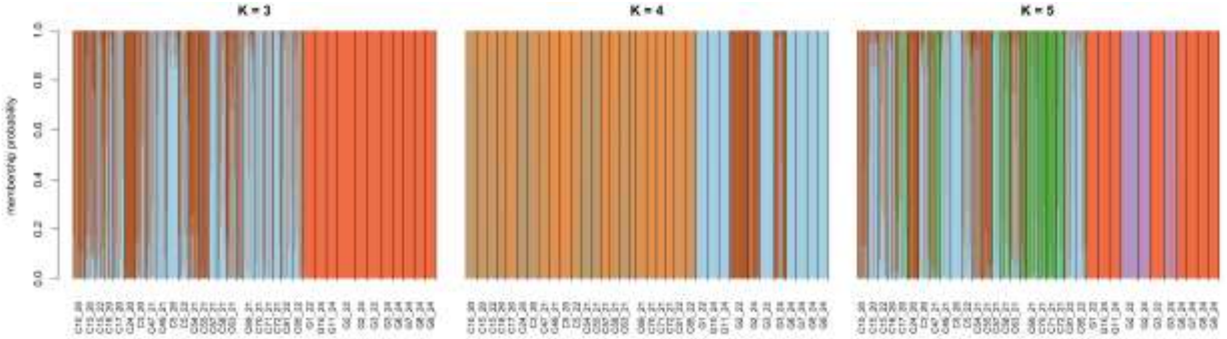


**Figure S1**: Genetic clustering of individuals inferred from Discriminant Analysis of Principal Components (DAPC) for K = 3 to K = 5. Each vertical bar represents a single individual, and colors denote the posterior assignment probability to each of the K genetic clusters inferred by k-means clustering.

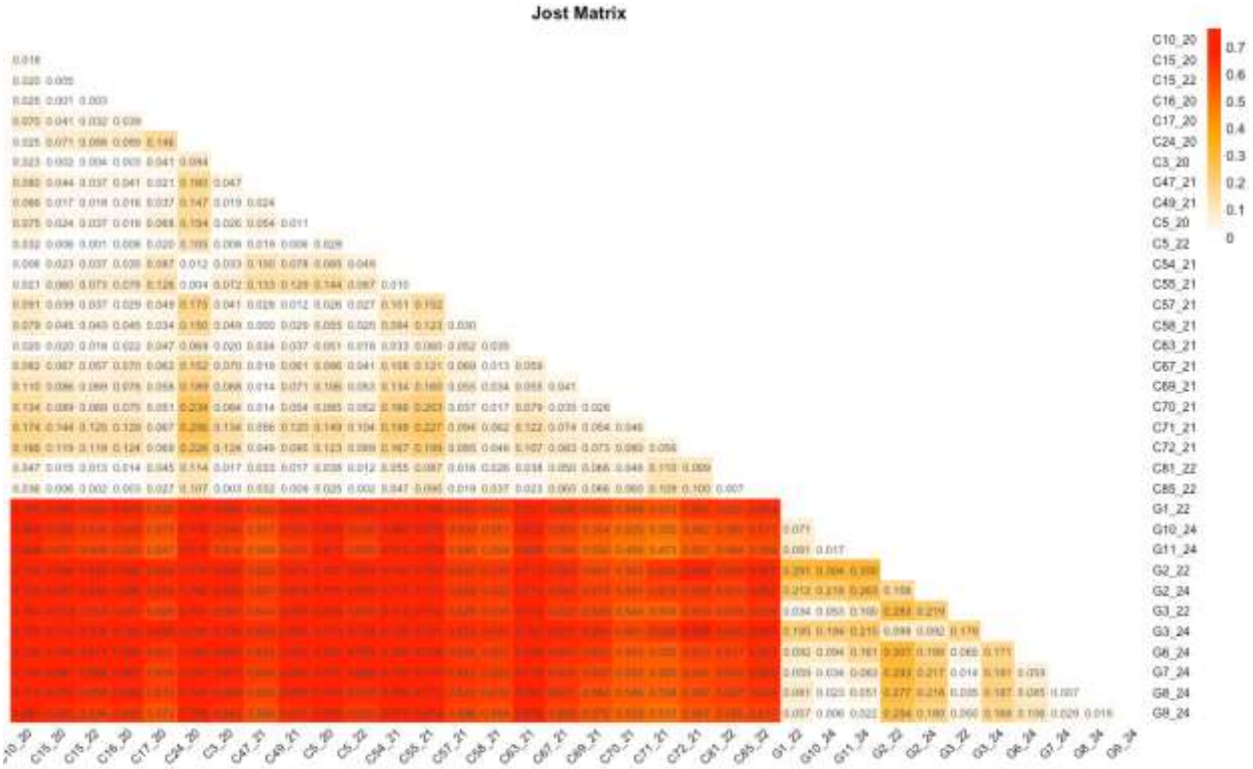

**Figure S2**: Heatmap displaying pairwise values of Jost's actual differentiation ($D_{est}$) between sampling sites across Coutada 9 (sites prefixed with "C") and Gorongosa National Park (sites prefixed with "G"). The color gradient ranges from white (low differentiation, approx 0) to intense red (high differentiation, $D_{est} > 0.7$).

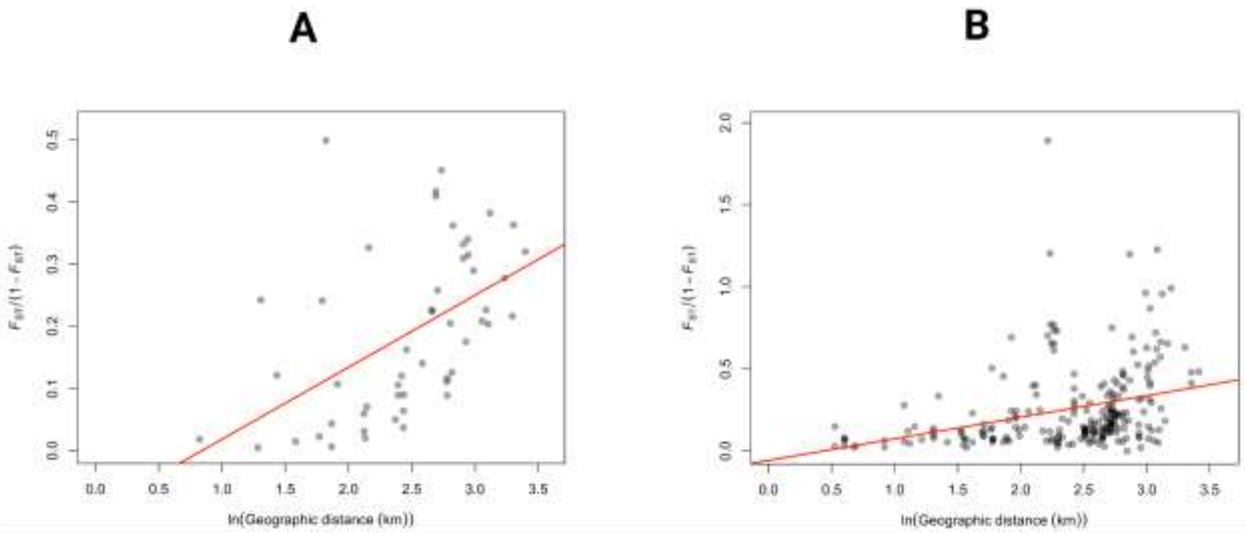


**Figure S3**: Patterns of isolation by distance (IBD) within conservation areas. Scatter plots showing linearized genetic differentiation ($F_{ST}$) as a function of the natural logarithm of geographic distance (ln(km)) between sampling sites in **(A)** Gorongosa National Park and **(B)** Coutada 9 Game Reserve. Pairwise $F_{ST}$ values were calculated following Weir and Cockerham (1984). Solid red lines represent the fitted ordinary least-squares linear regressions (y~x).

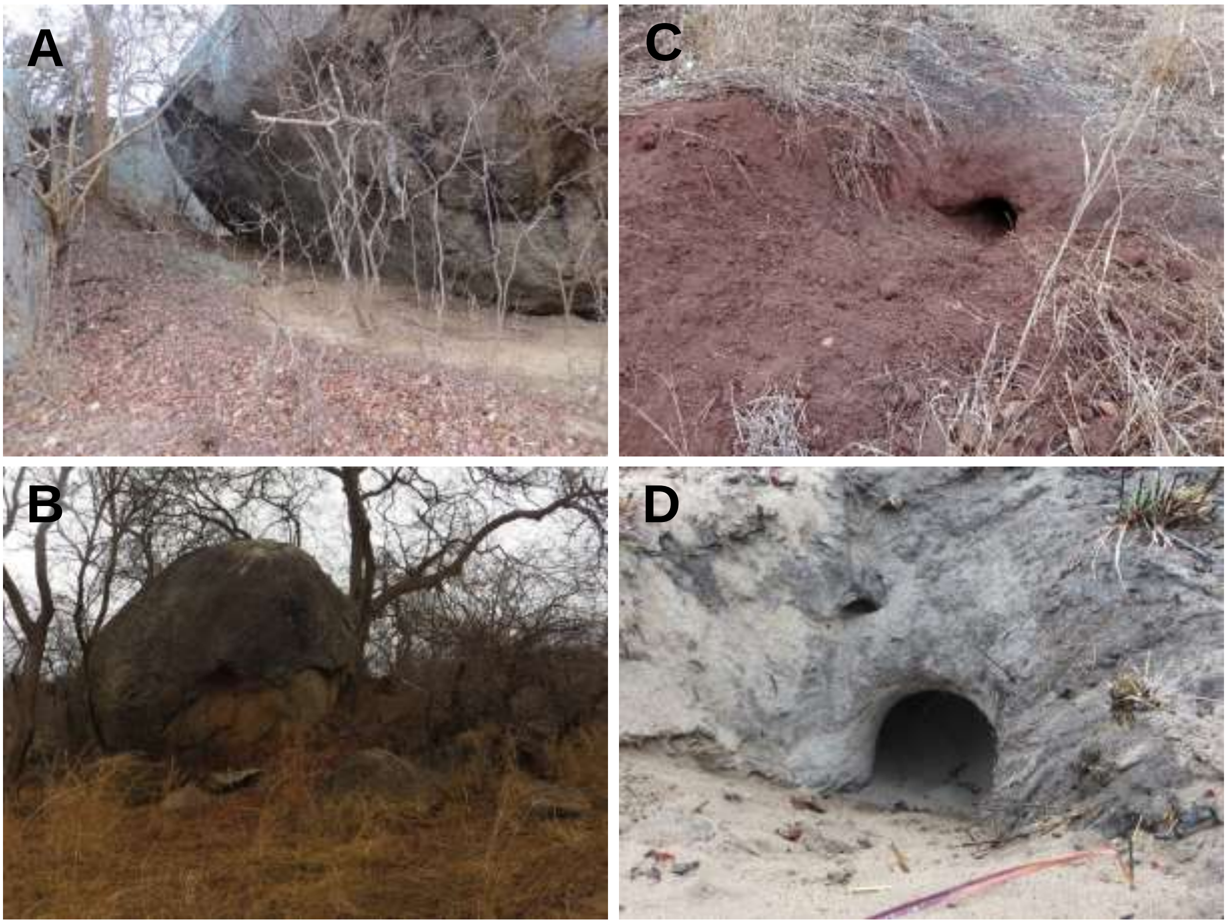


**Figure S4:** Pictures of two “rocks” sites and two “burrows” sites that were sampled for *Ornithodoros* ticks. **A**. and **B**. rocks. **C**. and **D**. burrows.

| | N_ind | N_alleles | $H_o$ | $H_e$ | $F_{IS}$ | HWE |
|---|---|---|---|---|---|---|
| **C10_20** | 30 | 1.563 | 0.106 | 0.137 | 0.264 | 0.750 |
| **C15_20** | 30 | 1.625 | 0.172 | 0.161 | 0.075 | 0.998 |
| **C15_22** | 30 | 1.875 | 0.194 | 0.202 | 0.131 | 0.303 |
| **C16_20** | 31 | 1.813 | 0.141 | 0.176 | 0.212 | 0.485 |
| **C17_20** | 30 | 1.938 | 0.278 | 0.282 | 0.059 | 0 |
| **C24_20** | 29 | 1.375 | 0.027 | 0.039 | 0.214 | 1 |
| **C3_20** | 29 | 1.750 | 0.153 | 0.180 | 0.192 | 0.938 |
| **C47_21** | 30 | 2.313 | 0.305 | 0.346 | 0.070 | 0.238 |
| **C49_21** | 30 | 1.688 | 0.159 | 0.171 | 0.044 | 0.999 |
| **C5_20** | 30 | 1.375 | 0.201 | 0.139 | -0.433 | 0.999 |
| **C5_22** | 29 | 2.063 | 0.213 | 0.208 | 0.027 | 0.873 |
| **C54_21** | 30 | 1.313 | 0.083 | 0.108 | 0.342 | 0.150 |
| **C55_21** | 30 | 1.438 | 0.077 | 0.105 | 0.337 | 0.848 |
| **C57_21** | 31 | 2.188 | 0.193 | 0.212 | 0.142 | 0.489 |
| **C58_21** | 30 | 2.250 | 0.372 | 0.362 | -0.003 | 0.708 |
| **C63_21** | 29 | 1.875 | 0.201 | 0.238 | 0.170 | 0.062 |
| **C67_21** | 30 | 2.125 | 0.195 | 0.310 | 0.269 | 0.254 |
| **C69_21** | 29 | 2.375 | 0.353 | 0.364 | 0.029 | 0.084 |
| **C70_21** | 29 | 2.375 | 0.304 | 0.376 | 0.155 | 0.002 |
| **C71_21** | 30 | 2.313 | 0.338 | 0.397 | 0.142 | 0.012 |
| **C72_21** | 24 | 2.375 | 0.392 | 0.369 | -0.012 | 0.010 |
| **C81_22** | 29 | 2.125 | 0.215 | 0.223 | 0.111 | 0.106 |
| **C85_22** | 30 | 1.938 | 0.164 | 0.196 | 0.178 | 0.079 |
| **G1_22** | 44 | 2.813 | 0.331 | 0.366 | 0.083 | 0.001 |
| **G10_24** | 29 | 3.500 | 0.445 | 0.507 | 0.091 | 0 |
| **G11_24** | 30 | 3.563 | 0.483 | 0.526 | 0.067 | 0.100 |
| **G2_22** | 51 | 2.688 | 0.389 | 0.422 | 0.053 | 0.276 |
| **G2_24** | 30 | 2.625 | 0.360 | 0.391 | 0.059 | 0.014 |
| **G3_22** | 42 | 3.000 | 0.366 | 0.402 | 0.057 | 0.032 |
| **G3_24** | 30 | 2.813 | 0.479 | 0.479 | -0.001 | 0.001 |
| **G6_24** | 26 | 2.688 | 0.276 | 0.350 | 0.227 | 0.008 |
| **G7_24** | 30 | 3.188 | 0.382 | 0.437 | 0.085 | 0.001 |
| **G8_24** | 27 | 3.313 | 0.435 | 0.484 | 0.094 | 0 |
| **G9_24** | 29 | 3.313 | 0.460 | 0.507 | 0.072 | 0 |

**Table S1**: Basic genetic diversity estimates for each studied population in Coutada 9 (sites prefixed with "C") and Gorongosa National Park (sites prefixed with "G") and. N_ind: number of individuals analysed. N_alleles: total number of alleles. $H_o$/ $H_e$: observed/expected heterozygosity. $F_{IS}$: intrapopulation fixation index. HWE: Global Hardy–Weinberg equilibrium deviations across all polymorphic loci within each population were assessed using Fisher's combined probability tests (p -value)

| Model | Model specification | Source of variation | df | Sum of squares | Mean squares | Variance component (σ) | Variation (%) | Φ statistic |
|---|---|---|---|---|---|---|---|---|
| **Model 1** | ~ Parc / Pop | Among parks | 1 | 4340.336 | 4340.336 | 4.923 | 51.16 | ΦCT = 0.512 |
| **Model 1** | ~ Parc / Pop | Among populations within parks | 32 | 1498.809 | 46.838 | 0.773 | 8.03 | ΦSC = 0.165 |
| **Model 1** | ~ Parc / Pop | Among individuals within populations | 906 | 3852.613 | 4.252 | 0.326 | 3.39 | ΦIS = 0.083 |
| **Model 1** | ~ Parc / Pop | Within individuals | 940 | 3383.717 | 3.600 | 3.600 | 37.41 | |
| **Model 1** | ~ Parc / Pop | Total | 1879 | 13075.475 | 6.959 | 9.622 | 100.00 | ΦIT = 0.626 |
| **Model 2** | ~ Year / Pop | Among years | 3 | 3345.098 | 1115.033 | 2.204 | 29.14 | ΦCT = 0.291 |
| **Model 2** | ~ Year / Pop | Among populations within years | 30 | 2494.048 | 83.135 | 1.434 | 18.96 | ΦSC = 0.268 |
| **Model 2** | ~ Year / Pop | Among individuals within populations | 906 | 3852.613 | 4.252 | 0.326 | 4.31 | ΦIS = 0.083 |
| **Model 2** | ~ Year / Pop | Within individuals | 940 | 3383.717 | 3.600 | 3.600 | 47.59 | |
| **Model 2** | ~ Year / Pop | Total | 1879 | 13075.475 | 6.959 | 7.564 | 100.00 | ΦIT = 0.524 |
| **Model 3** | ~ TypeOfSite / Pop | Among habitat types | 1 | 285.347 | 285.347 | 0.133 | 1.87 | ΦCT = 0.019 |
| **Model 3** | ~ TypeOfSite / Pop | Among populations within habitat types | 32 | 5553.798 | 173.556 | 3.068 | 43.05 | ΦSC = 0.439 |
| **Model 3** | ~ TypeOfSite / Pop | Among individuals within populations | 906 | 3852.613 | 4.252 | 0.326 | 4.58 | ΦIS = 0.083 |
| **Model 3** | ~ TypeOfSite / Pop | Within individuals | 940 | 3383.717 | 3.600 | 3.600 | 50.50 | |
| **Model 3** | ~ TypeOfSite / Pop | Total | 1879 | 13075.475 | 6.959 | 7.128 | 100.00 | ΦIT = 0.495 |

**Table S2:** Hierarchical AMOVA summary testing the effect of conservation area, sampling year, and habitat type on genetic diversity. Abbreviations: df, degrees of freedom; SS, sum of squares; MS, mean squares; σ, variance component; %, percentage of total variance. Hierarchical Φ -statistics follow Excoffier et al. (1992): ΦCT (among groups / total), ΦSC (among populations / within groups), ΦIS (among individuals / within populations), and ΦIT (individuals / total).

| Population | Co-ancestry method | 95% CI | Linkage disequilibrium method | 95% CI |
|---|---|---|---|---|
| **C10_20** | 1.2 | [0.9; 1.6] | 1.2 | [0.6; 2.2] |
| **C15_20** | 5.2 | [0.9; 13.3] | 2.5 | [1.1; 9.8] |
| **C15_22** | 2.4 | [1.2; 4.0] | 6.0 | [2.1; 19.2] |
| **C16_20** | 2.0 | [0.9; 3.5] | 5.1 | [1.5; 30.3] |
| **C17_20** | 2.6 | [0.9; 5.0] | 3.0 | [1.8; 8.0] |
| **C24_20** | 0.6 | [0.5; 0.6] | 0.4 | [0.2; 0.8] |
| **C3_20** | 6.9 | [0.5; 21.6] | 2.5 | [1.5; 6.8] |
| **C47_21** | 6.9 | [0.8; 19.3] | 5.5 | [2.7; 10.2] |
| **C49_21** | 1.4 | [1.1; 1.8] | 2.6 | [0.8; 20.3] |
| **C5_20** | ∞ | [∞; ∞] | ∞ | [0.0; ∞] |
| **C5_22** | 1.8 | [1.0; 3.0] | 3.0 | [1.5; 11.6] |
| **C54_21** | 1.0 | [0.8; 1.2] | 0.3 | [0.3; 0.3] |
| **C55_21** | 0.8 | [0.6; 1.1] | 1.4 | [0.5; 4.9] |
| **C57_21** | 19.5 | [0.0; 97.8] | 27.3 | [6.1; ∞] |
| **C58_21** | 6.9 | [1.2; 17.8] | 28.5 | [5.5; ∞] |
| **C63_21** | 9.8 | [0.2; 36.2] | 7.1 | [2.3; 22.8] |
| **C67_21** | 7.2 | [2.0; 15.9] | 20.1 | [4.7; ∞] |
| **C69_21** | 3.9 | [2.8; 5.2] | 13.2 | [3.3; 73.4] |
| **C70_21** | 2.3 | [1.1; 4.0] | 4.1 | [2.2; 9.8] |
| **C71_21** | 13.6 | [0.0; 68.3] | 10.0 | [2.8; 45.9] |
| **C72_21** | 6.9 | [0.0; 34.9] | 3.1 | [1.9; 8.9] |
| **C81_22** | 2.0 | [1.3; 2.9] | 8.2 | [2.9; 19.4] |
| **C85_22** | 3.2 | [0.7; 7.8] | 7.5 | [2.7; 18.4] |
| **G1_22** | 11.6 | [0.3; 42.6] | 15.0 | [5.0; 77.2] |
| **G10_24** | ∞ | [∞; ∞] | 271.8 | [46.3; ∞] |
| **G11_24** | ∞ | [∞; ∞] | 347.8 | [36.7; ∞] |
| **G2_22** | 1.7 | [0.9; 2.7] | 3.9 | [3.0; 7.5] |
| **G2_24** | 5.0 | [1.4; 11.0] | 2.8 | [2.0; 5.3] |
| **G3_22** | 41.3 | [0.0; 207.4] | 293.2 | [50.7; ∞] |
| **G3_24** | 4.4 | [2.6; 6.6] | 2.7 | [1.7; 6.7] |
| **G6_24** | 2.8 | [2.1; 3.6] | 3.3 | [2.0; 9.2] |
| **G7_24** | ∞ | [∞; ∞] | ∞ | [55.4; ∞] |
| **G8_24** | 10.8 | [1.8; 27.7] | ∞ | [47.0; ∞] |
| **G9_24** | ∞ | [∞; ∞] | 83.7 | [27.9; ∞] |

Table S3: Contemporary effective population size ($N_e$) estimates per sampled population. Estimates were computed using NEESTIMATOR v2.1 based on the molecular co-ancestry method (Nomura, 2008) and the linkage disequilibrium method (Waples & Do, 2008).

| Population | IAM | TPM | SMM |
|---|---|---|---|
| C10_29 | 0.187500 | 0.343750 | 0.593750 |
| C15_20 | 0.156250 | 0.230469 | 0.273438 |
| C15_22 | **0.024414** | 0.150391 | 0.248047 |
| C16_20 | 0.384766 | 0.577148 | 0.753906 |
| C17_20 | **0.006104** | **0.010498** | **0.033691** |
| C24_20 | 0.984375 | 0.984375 | 0.984375 |
| C3_20 | **0.037109** | 0.273438 | 0.421875 |
| C47_21 | **0.024719** | 0.076538 | 0.147888 |
| C49_21 | 0.101563 | 0.285156 | 0.544922 |
| C5_20 | **0.031250** | **0.031250** | **0.031250** |
| C5_22 | 0.473022 | 0.706055 | 0.904541 |
| C54_21 | **0.031250** | **0.031250** | **0.031250** |
| C55_21 | 0.281250 | 0.421875 | 0.578125 |
| C57_21 | 0.661377 | 0.866943 | 0.973877 |
| C58_21 | **0.001312** | **0.006714** | **0.024719** |
| C63_21 | **0.012207** | 0.080078 | 0.347656 |
| C67_21 | 0.054932 | 0.152710 | 0.270874 |
| C69_21 | **0.017670** | 0.060272 | 0.084412 |
| C70_21 | **0.009033** | 0.067688 | 0.151398 |
| C71_21 | **0.000427** | **0.010132** | **0.028992** |
| C72_21 | **0.012787** | 0.103882 | 0.280701 |
| C81_22 | 0.288574 | 0.617676 | 0.740234 |
| C85_22 | 0.367676 | 0.657593 | 0.792847 |
| G1_22 | 0.086304 | 0.195496 | 0.572388 |
| G10_24 | **0.002686** | **0.045288** | 0.213135 |
| G11_24 | **0.002686** | **0.033813** | 0.231567 |
| G2_22 | **0.002136** | **0.024719** | **0.045288** |
| G2_24 | **0.039246** | 0.147888 | 0.596130 |
| G3_22 | **0.045288** | 0.213135 | 0.312866 |
| G3_24 | **0.000153** | **0.000763** | **0.024719** |
| G6_24 | 0.101807 | 0.284668 | 0.765137 |
| G7_24 | **0.047119** | 0.108154 | 0.169800 |
| G8_24 | **0.000610** | **0.013306** | 0.136719 |
| G9_24 | **0.000427** | **0.001678** | 0.096863 |

**Table S4:** Signature of bottleneck (p-values from the one-tailed Wilcoxon signed-rank test for heterozygosity excess) for three different models of mutation in Coutada 9 Game Reserve and Gorongosa National Park. Sites indicated in red letters displayed significant signatures of bottleneck with the 3 models. Significant values ($p<0.05$) are indicated in bold.

| Locus name | Percentage of null allele (%) | Number of sites with null alleles | Stuttering | p-value for HW equilibrium |
|---|---|---|---|---|
| ms-2 | 3.8 | 1 | NO | [0.21 ; 0.24] |
| ms-24 | 1.7 | 0 | NO | [0.59 ; 0.68] |
| ms-30 | 0.1 | 0 | NO | [0.86 ; 0.94] |
| ms-35 | 1.6 | 0 | NO | [0.069 ; 0.090] |
| ms-48 | 1.4 | 0 | NO | [0.36 ; 0.42] |
| ms-59 | 0.5 | 0 | NO | [0.37 ; 0.54] |
| ms-61* | 2.1 | 3 | NO | <0.0001 |
| ms-63 | 3.4 | 0 | NO | [0.64 ; 0.71] |
| ms-71 | 0.6 | 0 | NO | [0.50 ; 0.53] |
| ms-73 | 0.9 | 1 | NO | [0.24 ; 0.30] |
| ms-76* | 12.5 | 14 | NO | <0.0001 |
| ms-78 | 4.4 | 2 | NO | [0.022 ; 0.028] |
| ms-81 | 1.1 | 0 | NO | [0.70 ; 0.74] |
| ms-87‡ | 3.2 | 2 | NO | [0.18 ; 0.22] |

| | | | | |
|---|---|---|---|---|
| ms-90 | 0.7 | 0 | NO | [0.60 ; 0.68] |
| ms-101 | 2.3 | 0 | NO | [0.42 ; 0.50] |
| ms-102 | 2.2 | 1 | NO | [0.18 ; 0.22] |
| ms-111* | 5.4 | 6 | YES | <0.0001 |
| ms-117 | 2.4 | 1 | NO | [0.73 ; 0.76] |

* These microsatellite loci were eliminated after quality evaluation. HW, deviation from Hardy Weinberg equilibrium tested with 10000 bootstraps. ‡ This microsatellite was not used for the second set of genotyping.

**Table S5**: Assessment of genotyping quality and Hardy–Weinberg equilibrium across the 19 microsatellite loci evaluated using Micro-Checker. For each locus, the table reports the estimated mean frequency of null alleles (percentage), the number of sampling sites with evidence of null alleles, the presence or absence of PCR stuttering artifacts, and the p -value (or confidence interval) for departure from Hardy–Weinberg (HW) equilibrium.

| Genotyping multiplex number | PCR multiplex (1:dilution) | Locus name | Fluorescent dye | Locus length (bp) | Primer concentration |
|---|---|---|---|---|---|
| Plex1 | PCR1 | ms-59 | FAM | 410 | 0.1µM |
| | | ms-35 | VIC | 79 | 0.15µM |
| | | ms-48 | NED | 246 | 0.1µM |
| | | ms-81 | NED | 448 | 0.3µM |
| | | ms-78 | PET | 291 | 0.2µM |
| Plex2 | PCR2 | ms-71 | FAM | 213 | 0.05µM |
| | | ms-102 | NED | 270 | 0.09µM |
| | | ms-24 | PET | 295 | 0.11µM |
| | | ms-63 | FAM | 412 | 0.3µM |
| | | ms-73 | NED | 456 | 0.25µM |
| Plex3 | PCR3 | ms-30 | FAM | 226 | 0.2µM |
| | | ms-101 | VIC | 159 | 0.35µM |
| | | ms-2 | NED | 286 | 0.45µM |
| | PCR4 | ms-117 | FAM | 438 | 0.2µM |
| | | ms-90 | NED | 457 | 0.35µM |

**Table S6:** Repartition of 15 microsatellite markers in three multiplexes for amplification and genotyping. PCR products were diluted by 1:20 before genotyping by capillary electrophoresis
